# Broadly inhibiting anti-neuraminidase monoclonal antibodies induced by trivalent influenza vaccine and H7N9 infection in humans

**DOI:** 10.1101/682450

**Authors:** Pramila Rijal, Bei Bei Wang, Tiong Kit Tan, Lisa Schimanski, Philipp Janesch, Tao Dong, John W. McCauley, Rodney S. Daniels, Alain R. Townsend, Kuan-Ying A. Huang

## Abstract

The majority of antibodies induced by influenza neuraminidase (NA), like those against hemagglutinin (HA), are relatively specific to viruses isolated within a limited time-window as seen in serological studies and the analysis of many murine monoclonal antibodies. We report three broadly reactive human monoclonal antibodies (mAbs) targeting N1 NA. Two were isolated from a young adult vaccinated with trivalent influenza vaccine (TIV), which inhibited N1 NA from viruses isolated from human over a period of a hundred years. The third antibody isolated from a child with acute mild H7N9 infection inhibited both group 1 N1 and group 2 N9 NAs. In addition, the antibodies cross-inhibited the N1 NAs of highly pathogenic avian H5N1 influenza viruses. These antibodies are protective in prophylaxis against seasonal H1N1 viruses in mice. This study demonstrates that human antibodies to N1 NA with exceptional cross-reactivity can be recalled by vaccination and highlights the importance of standardizing the NA antigen in seasonal vaccines to offer optimal protection.

**Importance:** Antibodies to the influenza NA can provide protection against influenza disease. Analysis of human antibodies to NA lags behind that for HA. We show that human monoclonal antibodies against NA induced by vaccination and infection can be very broadly reactive and able to inhibit a wide spectrum of N1 NAs between 1918 and 2018. This suggests that antibodies to NA may be a useful therapy, and that efficacy of influenza vaccines could be enhanced by ensuring appropriate content of NA antigen.

**Highlights of the paper:**

- Antibodies that inhibit influenza viruses with N1 neuraminidase (NA), with broad reactivity for viruses isolated between 1918-2018, can be isolated from human recipients of seasonal influenza vaccine
- Antibodies targeting N1 NA of human seasonal H1N1 viruses can cross-react with a variety of avian N1 neuraminidases
- Acute H7N9 infection can recall memory B cells to N1 NA and elicit cross-reactive antibodies to the group 1 N1 and group 2 N9 NAs
- Antibodies to N1 NA with this broad reactivity protect against lethal virus challenge

## Background

H1N1 virus entered the human population from birds in 1918. It is thought to have jumped from humans to pigs in that epoch, and it was from the pig that influenza virus was first isolated in 1931 (Shope, 1931) and shortly after from humans in 1933 through infection of ferrets (Smith W, 1933). H1N1 viruses circulated continuously in humans until 1958, when newly emerged H2N2 viruses replaced them. H1N1 virus reappeared in 1977 and continued to circulate until 2009. During this whole period, it underwent independent but continuous genetic and antigenic drift in humans and pigs. In 2009, a novel swine-origin H1N1 virus re-entered the human population and caused a pandemic. The accumulated sequence disparity between these independent descendants of the 1918 H1N1 virus had resulted in sufficient loss of cross-immunity to render most humans susceptible to infection by the porcine H1N1 virus.

Antibodies to the hemagglutinin (HA) and neuraminidase (NA) can independently provide protection from influenza disease (Couch et al., 2013, Monto et al., 2015, Memoli et al., 2016).The study of antibodies targeting NA has been under the shadow of those against HA, although there exists an extensive amount of evidence in favour of the protective immunity against NA. Previous work by Schulman, Webster, Kilbourne and colleagues showed the protective effects of anti-NA antibodies in mice and ferrets. Mice inoculated with virus or purified NA protein elicited protective immunity against NA (Schulman et al., 1968, Kilbourne et al., 2004). The anti-NA antibodies were shown to inhibit NA activity *in vitro* and reduce virus plaque size (Kilbourne et al., 1968b). Anti-NA immunity protected mice from infection, presumably by abrogating the release of virus from infected cells. Many groups subsequently elaborated the protective effects of antibodies against NA in animal models (Rockman et al., 2013, Doyle et al., 2013, Wan et al., 2013) [Reviewed by (Wohlbold and Krammer, 2014, Krammer et al., 2018, Eichelberger and Monto, 2019)].

Kilbourne and colleagues also showed that protective anti-NA antibodies are elicited in humans following natural infection (Kilbourne et al., 1968a, Schild, 1969) and exposure to inactivated whole-virus vaccine (Couch et al., 1974). Current challenge studies in humans also confirm the independent protective effect of antibodies against NA (Memoli et al., 2016). Finally, several groups have recently established the anti-NA antibody titre in human sera to be a correlate of protection in large clinical trials (Couch et al., 2013, Monto et al., 2015, Memoli et al., 2016).

Compared to a considerable literature on human mAbs against HA, the majority of mAbs targeting NA described to date are from mouse and rabbits which show relatively limited cross-reactivity. Among the first murine mAbs against NA - NC10 and NC41, specific to the N9 NA, were analysed for functional and structural characteristics (Malby et al., 1994, Lee and Air, 2002). The murine antibody CD6, which was protective against a limited range of N1 subtype viruses including seasonal H1N1, H1N1pdm09 and avian H5N1, was found to make several contacts with adjacent NA monomers. However, this antigenic epitope underwent amino acid substitution in recent seasonal H1N1 (post 2014) viruses and prevented CD6 binding (Wan et al., 2013, Wan et al., 2015).

Antibodies against NA act mainly through steric hindrance to block interaction of the active site of the enzyme with sialic acid templates, but may also invoke Fc-dependent protective mechanisms *in vivo* (Hashimoto et al., 1983, DiLillo et al., 2016, Jegaskanda et al., 2017). Antibody HCA-2, which was induced in rabbits by immunisation with a 9-mer conserved peptide from the NA active site (residues 222-230), is known to bind to the active site (Doyle et al., 2013). This antibody reacts in western blots with a very wide range of NAs, and cross-inhibits multiple strains and subtypes from influenza A and B subtypes, but only at high concentration. HCA-2 offers only partial protection, even at the high antibody dose of 60 mg/kg, and can be affected by amino acid substitutions in the active site that lead to reduced susceptibility to NA inhibitors (Doyle et al., 2013). The requirement for such a high concentration of HCA-2 is probably because it reacts with a linear epitope exposed predominantly after denaturation of NA. Thus, there is scope for potent and broadly reactive human mAbs against NA that confer better protection and could be used therapeutically.

Owing to high sequence diversity in the globular head of HA, humans produced broadly reactive antibodies to the conserved stalk of HA after exposure to H1N1pdm09 virus, targeting shared epitopes in the stalks of earlier seasonal H1N1 and H1N1pdm09 viruses. (Wrammert et al., 2011, Li et al., 2012). Antibodies against NA are less well studied in this context, but recently broadly reactive anti-NA antibodies have been isolated from humans after H1N1pdm09 virus infection (Chen et al., 2018). The NA of H1N1pdm09 viruses may have reactivated B cell memory for rare epitopes shared with the N1 of earlier human seasonal viruses. The authors could isolate such antibodies only after natural infection, not after vaccination. They confirmed that the NA antigen is poorly represented in many sub-unit vaccines, and that the quality and quantity of NA in different vaccines varies [Reviewed by (Marcelin et al., 2012)].

Despite this variability, we report a panel of anti-NA mAbs with exceptional broad reactivity, isolated from human donors after influenza vaccination or infection. Two broadly reactive human mAbs to N1 NA, isolated from a vaccinated individual inhibited the enzymatic activity of N1 NAs from viruses circulating in the course of the last 100 years. In addition, both mAbs cross-inhibited many N1 NAs from highly pathogenic avian influenza H5N1 viruses. The antibodies were effective prophylactics protecting mice against the highly lethal Cambridge variant of H1N1 virus A/PR/8/1934, and in the highly sensitive DBA/2 mouse strain challenged with a H1N1pdm09 virus. We also describe an antibody induced by acute H7N9 infection that cross-reacts between the human seasonal and avian N1 (group 1) and avian N9 (group 2) NAs. These exceptionally broadly reactive anti-NA mAbs may be found rarely but they offer the hope of developing vaccines that could induce them.

## Results

### Anti-neuraminidase mAbs from human donors

Two antibodies AG7C and AF9C were isolated from an adult (aged 23; donor C) vaccinated with 2014/15 northern hemisphere TIV containing A/California/7/2009 (Reassortant NYMC X-179A) (H1N1); A/Texas/50/2012 (Reassortant NYMC X-223) (H3N2), B/Massachusetts/2/2012 (Reassortant NYMC BX-51B); all at 15 µg/0.5 mL (AdimFlu-S produced by Addimmune Corporation, Taiwan) (Table 1). A third antibody, Z2B3, was isolated from a Chinese male child (donor Z) with a mild H7N9 infection in 2013; two more antibodies Z2C2 and Z1A11 were isolated from this donor. Similarly, three more N9 mAbs were isolated from donors W and K who were hospitalised with H7N9 virus infection (Table 1). Antibodies to H7 HA from donors Z and K are already reported (Huang et al., 2018).

**Table 1.** List of donors and anti-NA antibodies isolated

| Donor | Age, Gender & Collection year | Antigen exposure | Antibodies isolated | mAbs Specificity |
| --- | --- | --- | --- | --- |
| Donor C | 23, Male, 2014 | 2014/15 inactivated TIV (AdimFlu-S) | AG7C, AF9C | Specific to N1 NA |
| Donor Z | 6, Male, 2013 | Mild H7N9 infection | Z2B3, Z2C2, Z1A11 | Cross-reactive to N1 and N9 NAs |
| Donor W | 7, Female, 2013 | Severe H7N9 infection 2013 | W1C7 | N9 NA |
| Donor K | 39, Male, 2014 | Severe H7N9 infection 2013 | P17C, F4C | N9 NA, Weak N1 |

### Inhibitory breadth of anti-N1 NA mAbs against human H1N1 viruses

We focused our analysis on three mAbs: AG7C, AF9C and Z2B3 since other antibodies were either of limited specificity or weaker in inhibition of NA. These three mAbs were tested for the inhibition of NA activity of H1N1 viruses isolated between 1934 to 2018, in an Enzyme Linked Lectin Assay (ELLA) (Figures 1, 2), and for inhibition of the enzyme activity of the 1918 pandemic H1N1, and avian N1 and N9 NAs as recombinant proteins (Figure 3).

**Figure 1.**
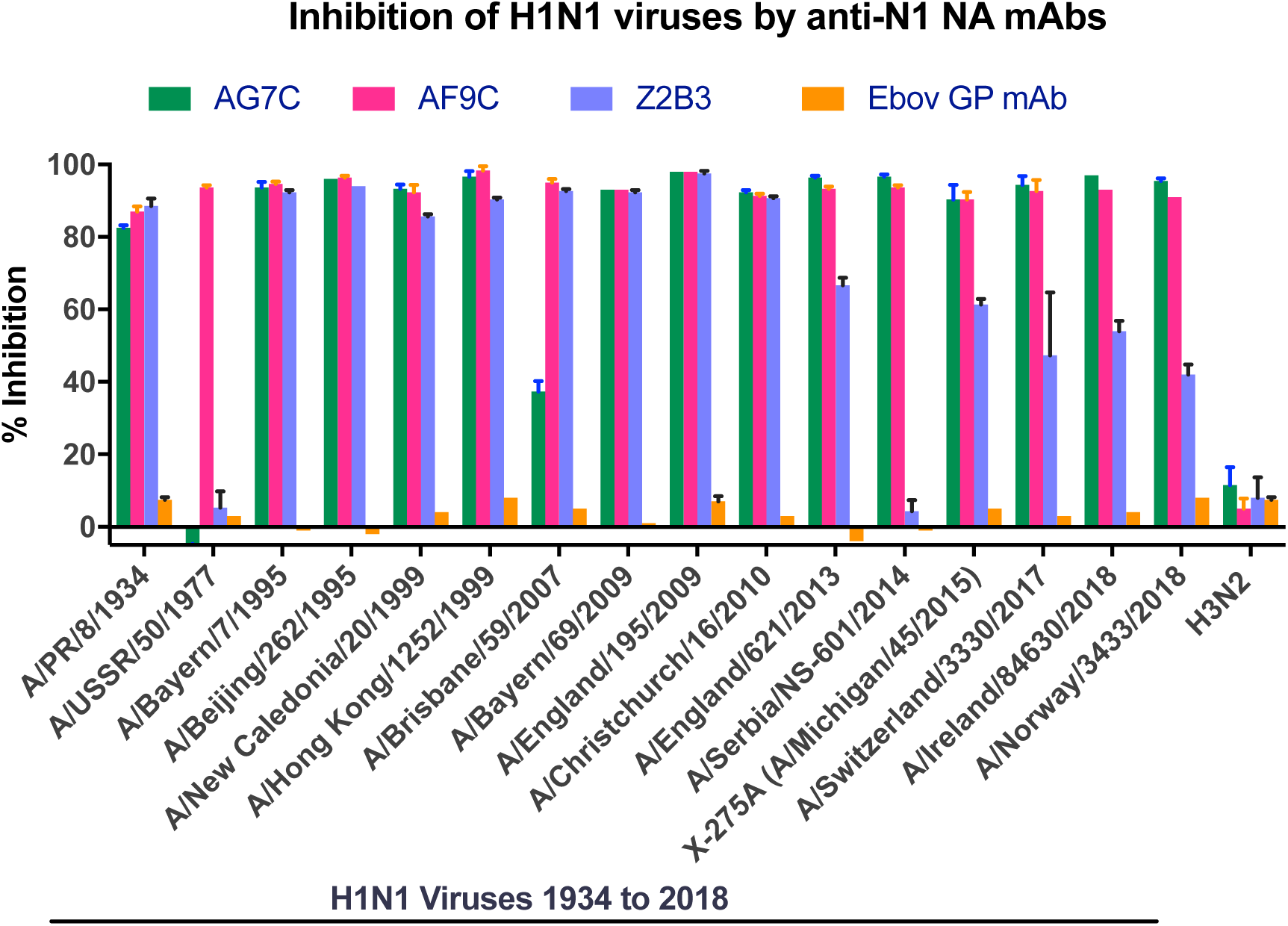
Inhibition of H1N1 viruses by mAbs targeting N1 NAs. Percentage inhibition of activity by mAbs, at 20 μg/ml, targeting N1 NAs are shown. H3N2 virus (X-31) was used as a negative control virus and a mAb targeting Ebolavirus glycoprotein was used as a negative control antibody.

**Figure 2.**
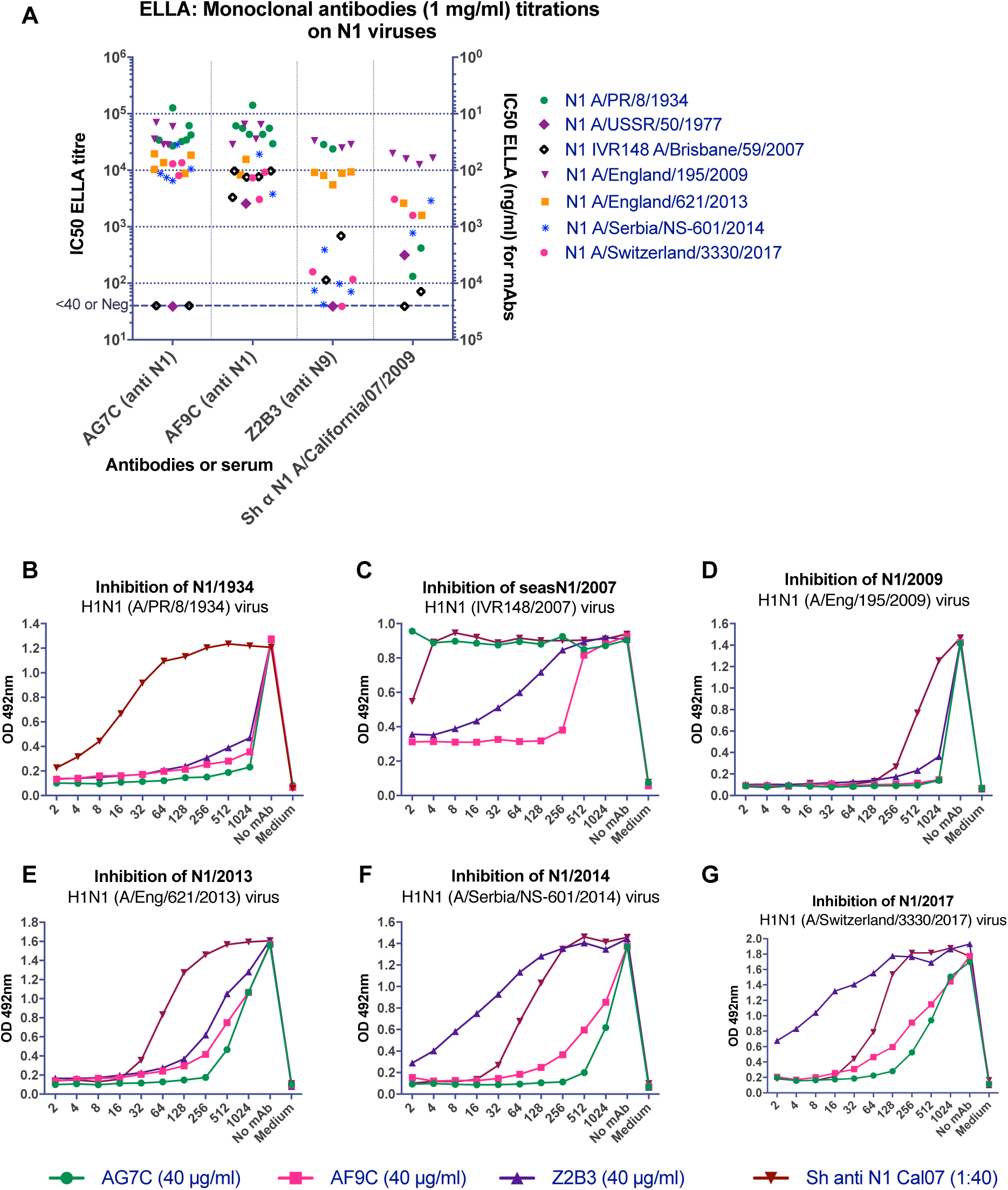
ELLA titrations of mAbs against selected H1N1 viruses. AG7C and AF9C are N1 NA-specific antibodies. Z2B3 is a N9 and N1 NA-cross-reactive antibody. Sheep anti-H1N1pdm09 N1 (A/California/07/2009) serum was used as a positive anti-N1 NA control. A) ELLA IC_50_ values of anti-N1 mAbs shown as titrating from 1 mg/ml on left Y-axis to compare with sheep sera and the 50% inhibiting concentration as ng/ml shown on right Y-axis. Each point represents an independent measurement. B-G) NA Inhibition curves for H1N1 viruses from year 1934 to 2017. Note the loss of titre of mAb Z2B3 on viruses isolated after 2014 (F, G). Experiments were done at least three times. Representative graphs are shown.

**Figure 3.**
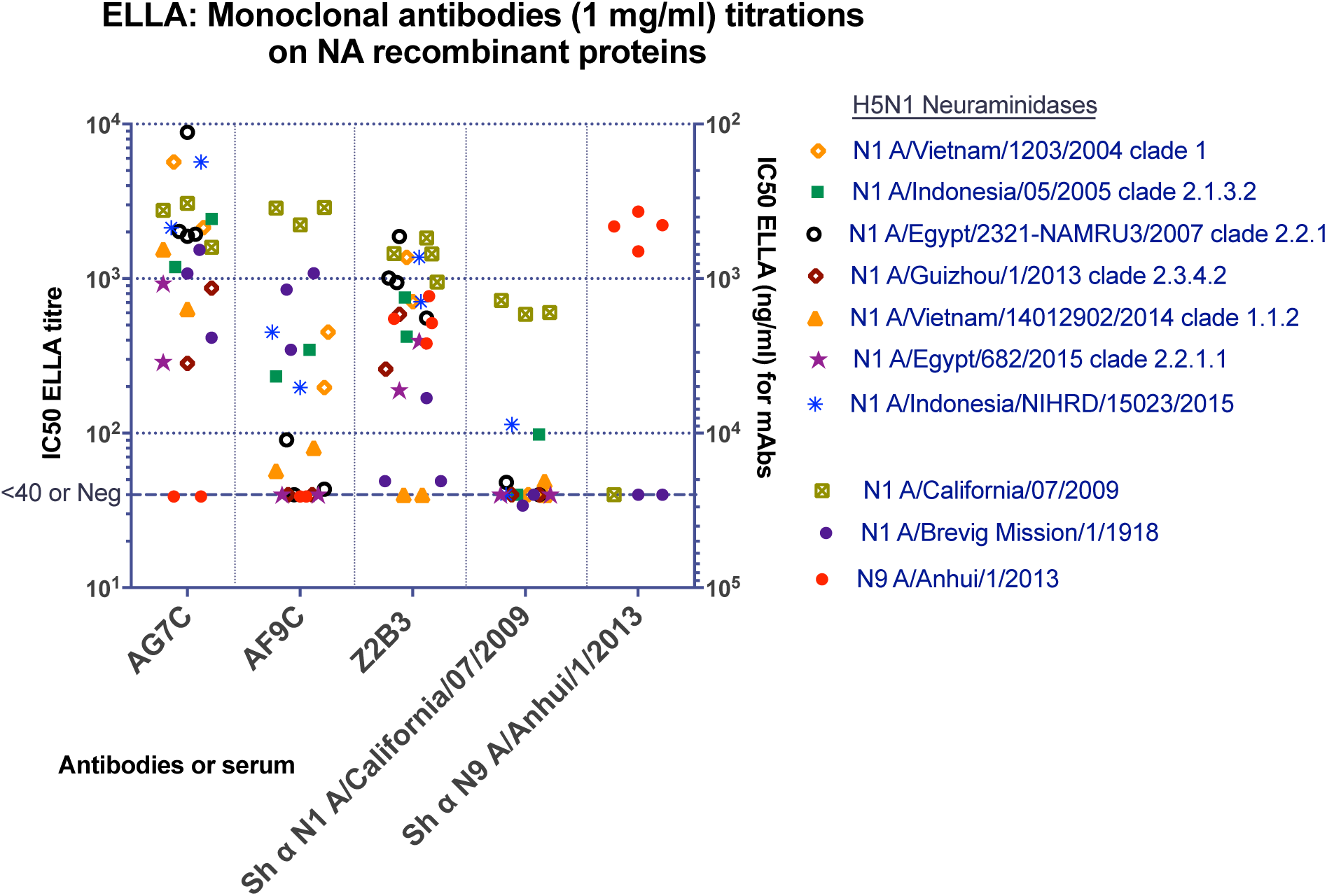
Antibodies (1mg/ml) titrated against recombinant NA proteins by ELLA. Sheep sera raised against H1N1pdm09 (A/California/07/2009) and H7N9 (A/Anhui/1/2013) viruses were used as controls. Each point represents an independent measurement.

The mAbs were titrated by ELLA and the concentrations required to give 50% inhibition (IC_50_) of NA activity were calculated by linear interpolation. The titres yielded by a 1 mg/ml solution were then calculated and plotted for comparisons to control hyper-immune sheep sera obtained from the National Institute for Biological Standards and Controls (NIBSC) (Figure 2, 3). On the secondary Y-axis, IC_50_ titres are shown in ng/ml.

AF9C inhibited the NA activities of all H1N1 viruses tested which represented the H1N1 viruses that have circulated in humans for over 100 years (Figures 1-3). AG7C showed a slightly different specificity in that it was weak or failed to inhibit the NAs from A/Brisbane/59/2007 and A/USSR/90/1977 (Figure 2). mAb Z2B3, cross-reactive with N9 NA, also showed a broad recognition of N1 NAs but was weak with A/Brisbane/59/2007 and failed to inhibit A/USSR/90/1977 NAs (Table 1; Figures 1, 2). Unlike AG7C and AF9C, Z2B3 had greatly reduced activity against recent clade 6B.1 H1N1pdm09 viruses isolated after 2014 (Figure 2).

Figure 2 shows that AG7C and AF9C titrate predominantly between 1:4,000 and 1:40,000 (IC_50_ ∼250-25 ng/ml) on the set of viruses shown, with the exception that AG7C fails to inhibit N1 NA form A/Brisbane/59/2007. By contrast Z2B3 gave similar titres on A/PR/8/1934, A/England/195/2009 and A/England/621/2013 but had drastically reduced titres on A/USSR/90/1977 and the representative recent clade 6B.1 H1N1pdm09 viruses A/Serbia/NS-601/2014 and A/Switzerland/3330/2017, indicating that the genetic and associated antigenic drift in these viruses had resulted in a major alteration in the epitope recognised by Z2B3. The control hyper-immune sheep serum to A/California/07/2009 N1 showed limited cross-reactivity on recently drifted or older (former seasonal) viruses with only weak activity against N1 NA from A/PR/8/1934. The sheep anti-H7N9 (A/Anhui/1/2013) serum contained anti-N9 NA antibodies that did not cross-react with any NAs expressed by these H1N1 viruses.

### The inhibitory activity of broadly reactive anti-N1 mAbs against NAs of avian H5N1 viruses

To avoid handling avian influenza viruses, we titrated the mAbs for inhibition of recombinant N1 NAs from a range of H5N1 viruses isolated from infected humans representing several HA-clades, from pandemic virus A/Brevig Mission/1/1918 and N9 NA from H7N9 virus A/Anhui/1/2013, produced in HEK293 cells, with N1 NA from A/California/07/2009 as a positive control (Figure 3).

AG7C inhibited all of the N1 NAs representing H5N1 viruses between 2004 and 2015 and the N1 NA from the 1918 pandemic virus A/Brevig Mission/1/1918. AF9C showed similar activity on the N1 NAs from A/California/07/2009 and A/Brevig Mission/1918 but was clearly weaker against the N1 NAs from H5N1 viruses. Neither AG7C nor AF9C inhibited the N9 NA. By contrast Z2B3 inhibited the H1N1pdm09 NA, the 2013 N9 NA and most of the avian N1 NAs at moderate IC_50_ values that were in general weaker than for mAb AG7C; it inhibited the 1918 N1 NA weakly The control hyper-immune sheep serum against H1N1pdm09 NA showed a titre >1:400 with A/California/7/2009 N1 NA, with minimal cross-reactivity with avian N1 NAs, 1918 N1 NA or the 2013 N9 NA. The control sheep serum against N9 NA inhibited N9 but not N1 NAs.

### Anti-N9 NA mAbs cross-reactive with N1 NA

Among six anti-N9 NA mAbs isolated from three donors exposed to H7N9 virus and tested by ELLA, three inhibited recombinant N9 NA (Figure 4). Two N9 NA-inhibiting mAbs were isolated from donor Z, where Z2B3 was a strong inhibitor and Z2C2 was a weak inhibitor (Figure 2A). All three mAbs from donor Z were cross-reactive with N1 NA (Figure 4C) and strongly inhibited the H1N1pdm09 (A/England/195/2009) N1 NA (Figure 4B). This suggests that 6-year old donor Z may have made a primary antibody response to the H1N1pdm09 N1 NA, and subsequent infection with H7N9 stimulated the memory B cells to an epitope conserved between N1 and N9 NAs. Notably, Z2B3 and Z2C2 have longer heavy chain CDR3 domains than other mAbs and although Z2B3 and AF9C are both encoded by the same VH gene (VH1-69), the CDR3 amino acid sequences are significantly different.

**Figure 4.**
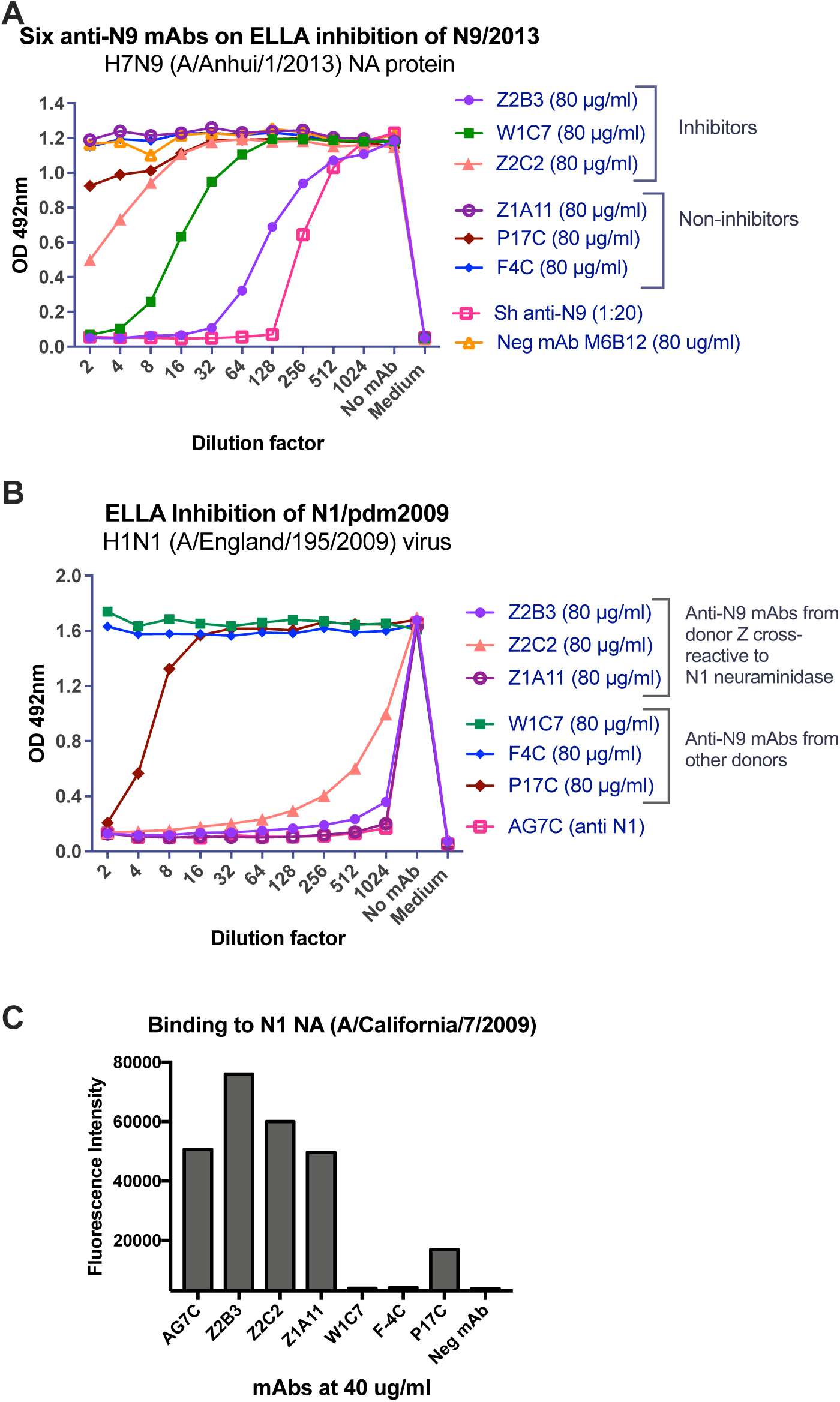
Inhibition of NA activity by mAbs isolated from donors exposed to H7N9 virus. A) ELLA activity of six anti-N9 antibodies on N9 NA (A/Anhui/1/2013). Sheep sera raised against H7N9 virus (A/Anhui/1/2013) acts as a positive control. Anti-N2 NA mAb M6B12 was used as a negative control. B) Cross-inhibition of N1 NA by some anti-N9 NA mAbs. Anti N1-NA mAb (AG7C) is a positive control. C) Binding of anti-N9 NA mAbs to H1N1 (X-179A A/California/7/2009) infected MDCK-SIAT cells. Experiments were performed at least twice, and representative graphs are shown.

Antibodies from donors W (W1C7) and K (P17C, F4C) were found to bind N9 NA in an indirect immunofluorescence screen (not shown). W1C7 and F4C were specific for N9 NA, and W1C7 had a weak inhibitory effect in ELLA on N9 (Figure 4). P17C cross-reacted with N1 NA with low level of binding and showed weak inhibition by ELLA (Figure 4B, C).

Antibodies from donor Z have higher numbers of amino acid substitutions in the variable regions of heavy and light chains, compared to those in mAbs from other donors (Table 2). The number of substitutions in VH of mAbs Z2B3, Z2C2 and Z1A11 are 8, 13 and 17 respectively, whereas there are none, 1 and 1 respectively in mAbs W1C7, P17C and F4C. This suggests the mAbs from donor Z are of memory B cell origin while those from donors W and K resulted from de-novo responses to acute H7N9 infection.

**Table 2.**
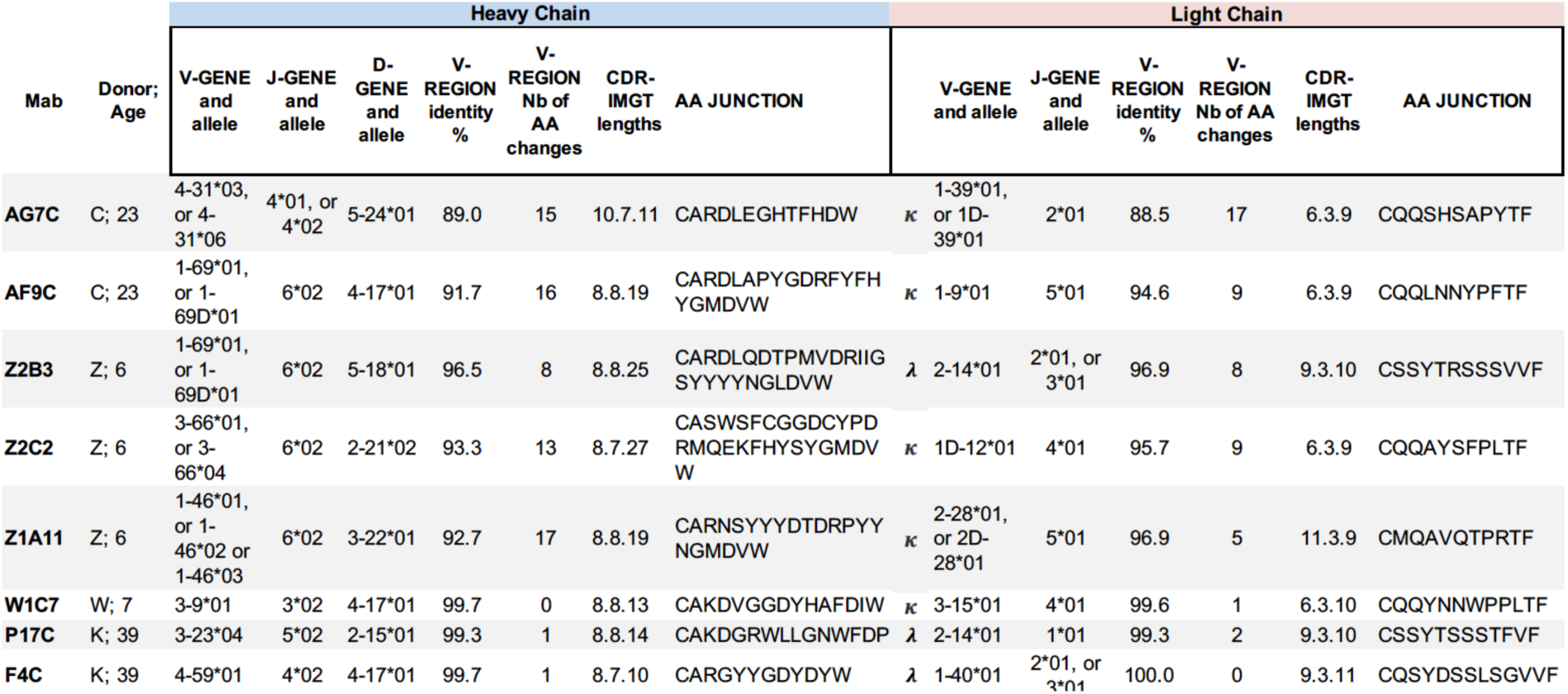
Encoding gene analysis of antibodies

### Anti-NA mAbs provide prophylactic protection *in vivo*

All three of the anti-N1 NA mAbs, AG7C, AF9C and Z2B3, protected 100% of mice from challenge with 10^4^ TCID_50_ of A/PR/8/1934 virus (equivalent to 1000 LD_50_) when given at a dose of 10 mg/kg 24 hours before infection (p<0.001; Figures 5A, B). They prevented any weight loss whereas mice that received an anti-N2 NA mAb (M6B12) succumbed to ≅20% weight loss by day 5 and were humanely culled. An antibody to the H1 stem T1-3B (Huang 2015) provided a positive control for protection.

**Figure 5.**
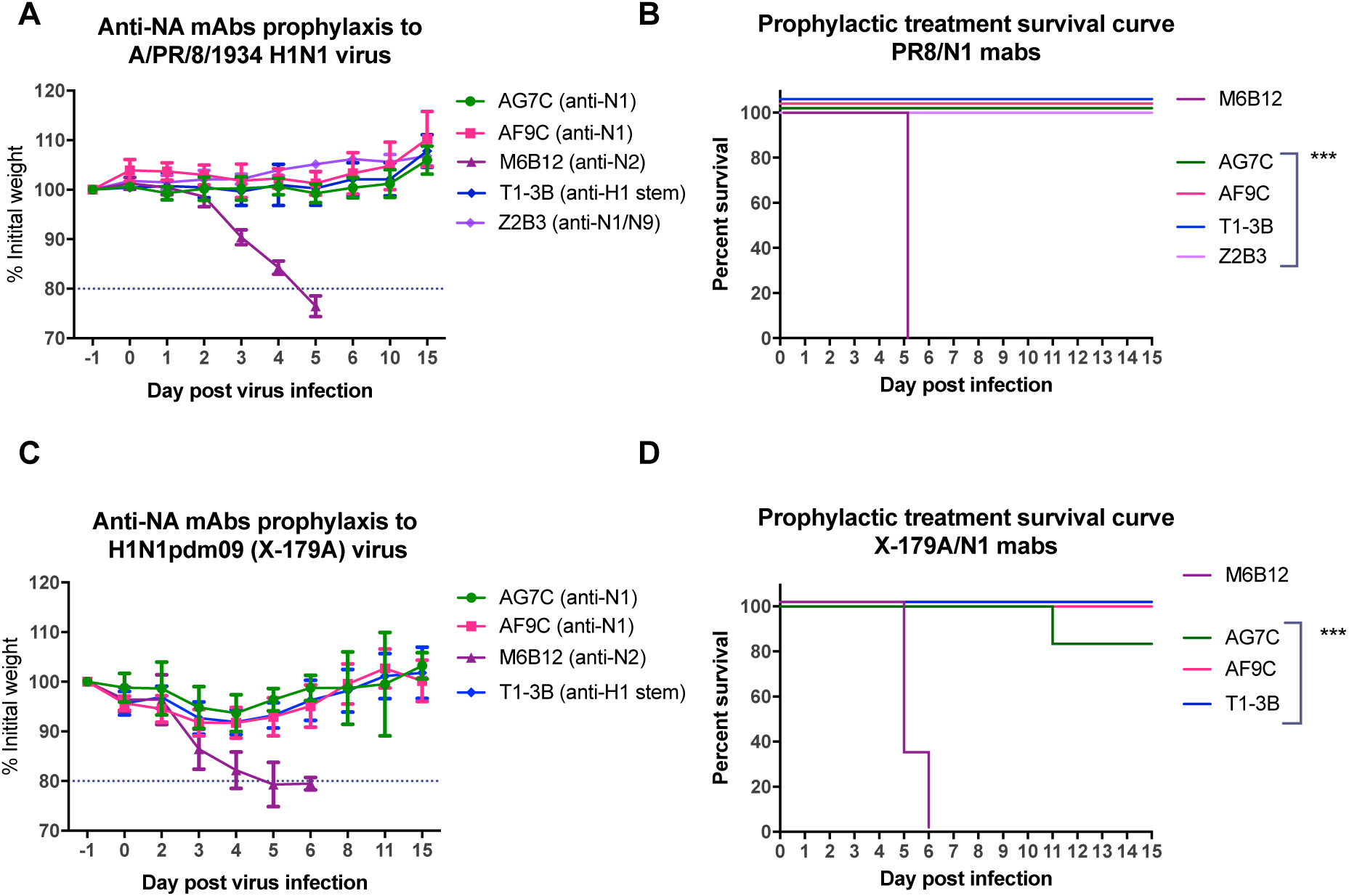
In vivo prophylactic protection by anti-N1 NA mAbs. Mice (n=6/group) were administered AG7C and AF9C mAbs at 10 mg/Kg. Weight loss following infection was measured and ≅20% loss was considered as the predefined endpoint. Anti-H1 HA mAb (T1-3B) cross-reactive with X-179A and A/PR/8/1934 viruses, is a positive control and an anti-N2 NA specific mAb (M6B12) a negative control. Experiments were performed at least twice and representative data from individual experiments are shown here. (A, B) Anti-N1 NA mAbs protect Balb/C female mice completely against 10^4^ TCID_50_ of A/PR/8/1934 virus, without any weight loss (p<0.001). (C,D) Anti-N1 NA mAbs protect DBA/2 female mice completely against a lethal dose (∼150 LD_50_) of X-179A virus (A/California/7/2009), with only 5-10% weight loss (p<0.001). One mouse treated with AG7C relapsed on day 7 and was culled after losing >20% weight; it is possible that a mAb-escape influenza variant may have emerged in this mouse.

In another experiment, DBA/2 mice, that are uniquely susceptible to influenza infection (Pica et al., 2011) were treated with AG7C and AF9C antibodies 24 h before infection with 10^4^ TCID_50_ of X-179A (equivalent to 150 LD_50_) virus, a reassortant containing the H1N1pdm09 vRNAs from A/California/07/2009 (Figures 5C,D).

Treated mice were protected from ≅20% weight loss (p<0.001), whereas mice receiving a non-specific antibody had to be culled on days 5 or 6. One out of 6 mice in the AG7C group was sacrificed on day 11 after losing >20% weight. In these prophylactic protection experiments, anti-NA mAbs were as protective as T1-3B, the positive control anti-HA stalk mAb (Huang et al., 2015).

## Discussion

We show in this paper that broadly reactive and protective antibodies to N1 NA can be isolated from vaccinated and infected individuals, presumably due to the conservation in surface structure between N1 NAs (Figure 6A). The two N1 subtype specific mAbs AG7C and AF9C were isolated from the same donor who had been vaccinated in 2014 with AdimFlu-S TIV in Taiwan. AG7C inhibits N1 NAs from H1N1 viruses isolated between 1918-2018. Although previous investigations of subunit vaccines have found varying and usually low levels of NA antigen (Chen et al., 2018, Krammer et al., 2018) in this case there was clearly enough to induce a response.

**Figure 6.**
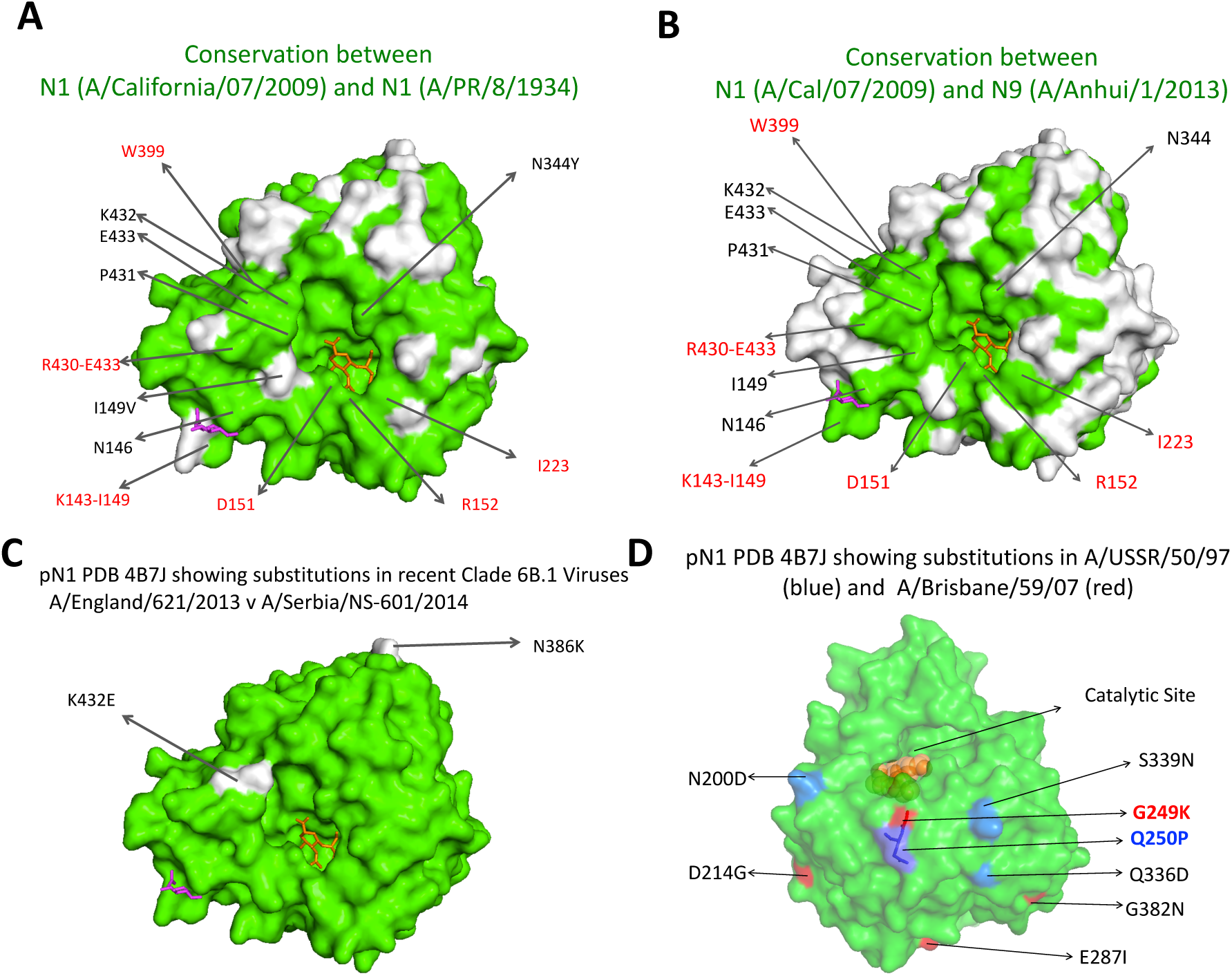
Comparisons of conserved and variable surface residues between NA subtypes. (A) Conserved molecular surface shown in green between H1N1pdm09 A/California/07/2009 and H1N1 A/PR/8/1934 (PDB 4B7J). (B) Conserved molecular surface shown in green between H1N1pdm09 (A/California/07/2009) and H7N9 (A/Anhui/1/2013) (PDB 4B7J). (C) Difference in molecular surface (shown in white) between H1N1pdm09 viruses A/England/621/2013 and A/Serbia/NS-601/2014 (D) Key amino acid substitutions in two H1N1 virus NAs that mAb AG7C inhibits poorly - A/USSR/50/1997 (shown in blue) and A/Brisbane/59/2007 (shown in red) compared to NA of H1N1pdm09. These amino acid positions were inferred from the NA sequences alignment in Figure S2. G249 and Q250P are likely to form part of the binding footprint of mAb AG7C.

The very broad reactivity of these mAbs with N1 NAs, covering the complete period of H1N1 virus circulation in humans, may have been induced by exposure to the significantly different NA derived from the H1N1pdm09 virus. Both mAbs show significant sequence divergence (Table 2) suggesting that they originated from a memory population which went through multiple rounds of selection in germinal centres following previous exposures to influenza. Both mAbs provided prophylactic protection in mice against the highly virulent variant of A/PR/8/1934 (the Cambridge strain) (Grimm et al., 2007) and, in ultra-sensitive DBA/2 mice, against infection with H1N1pdm09 X-179A (A/California/7/2009). In an earlier paper Chen et al. described similar anti-N1 NA antibodies that reacted with viruses spanning the period 1918-2009 (Chen et al., 2018).

The third antibody Z2B3 was isolated from a child who experienced a mild infection with H7N9 virus in 2013. It was unusual in being cross-reactive with group 1 (N1) and group 2 (N9) NAs. Two similar antibodies were isolated from this donor, both of which inhibited N1 NA with some level of cross-reaction with N9 NA (Figure 4), which we interpret to imply that they were selected from a subpopulation of memory cells induced previously by N1 NA. Examination of the structure of the N1 and N9 NAs reveals a region of conserved surface around and within the active site of the enzyme, as a possible binding site for Z2B3 (Figure 6B).

Mab Z2B3 showed good reactivity with the H1N1pdm09 virus A/England/621/2013, but poor reactivity with a later clade 6B virus, A/Serbia/NS-601/2014. These two viruses showed non-conservative amino acid substitutions of only N386K and K432E in the head of NA (Figure 6C). The former site is similarly substituted in the N9 NA that Z2B3 recognizes, which suggests that K432 is within the footprint of mAb Z2B3. K432 falls within a known epitope recognized by anti-N9 NA antibodies (Malby et al., 1994, Tulip et al., 1992). The crystal structure of a N9 NA-mAb complex, N9-NC10, involved a contact between D56 of the antibody H-chain and K432 of N9 NA (GRP**<u>K</u>**EDK; PDB 1NMB).

K432 was conserved prior to 2013 but underwent substitution in 2014, K432E, which became dominant thereafter. We suggest that N1 NA has been under strong evolutionary pressure from broadly cross-reactive antibodies induced by the H1N1pdm09 NA, that were selected from memory B cells raised against NA(s) of earlier virus(es). Just as the conserved stalk of HA has shown a capacity for evolution under pressure from antibody selection (Doud et al., 2018), the NA may similarly be forced to drift antigenically by broadly cross-reactive antibodies induced by the H1N1pdm09 viruses (Gao et al., 2019).

With this in mind we examined the region of the NA surface recognized by broadly reactive antibodies described by Chen et al. that inhibited or bound N1 NAs of viruses isolated between 1918-2009 but not clade 6B H1N1pdm09 viruses (Gao et al., 2019, Chen et al., 2018). Some of these antibodies lost binding to N1 NAs with substitutions in a set of site-specific mutants (Wan et al., 2015, Gao et al., 2019). Many of these antibodies also did not inhibit A/Brisbane/59/2007. mAb AG7C showed a similar reactivity profile and may have been affected by substitutions G249K and Q250P that are common to the non-reactive NAs. These residues are exposed on the periphery of the catalytic site (Figure 6D). The preceding residue N248 was substituted (N248D) in the H1N1pdm09 viruses isolated post 2009 and caused a loss of recognition by one of the antibodies described by Chen et al. However, this substitution is tolerated by mAb AG7C. There are rare natural isolates that have substituted these residues (G249E/R and Q250R) indicating that even the broadly reactive mAbs can be thwarted by virus antigenic drift. Further structural work to define the epitopes recognised by Z2B3, AG7C and AF9C is in progress.

It has become clear that exposure to viruses that differ significantly from those circulating, can select responses to epitopes in both HA and NA that are shared between the incoming virus and the seasonal viruses in circulation, derived from the memory B cell population (Henry et al., 2018). While antibodies against new epitopes can also be generated, even in the elderly (Huang et al., 2018), it appears that they are initially at a disadvantage but may overtake and become dominant with time (Lee et al., 2019, Henry et al., 2019). It is these high affinity and relatively specific antibodies that are mainly detected in serological surveys (Fonville et al., 2014). It would be wise to assume that all of these epitopes, both new and conserved, can drift under pressure from antibody selection. The inevitable implication is that updating influenza vaccines may have to continue but broadening the memory B cell population by vaccination with as wide a range of groups 1 and 2 HAs and NAs as possible might be a logical way of preparing the ground for a strong response to an unknown future pandemic virus.

## Materials and Methods

### Media, Reagents and Tissue Culture

MDCK-SIAT1 cells and adherent 293T cells (ECACC) were grown in D10 - DMEM medium (Sigma D5796) supplemented with 10% v/v foetal calf serum (Sigma F9665), 2 mM glutamine, 100 U/mL penicillin and 100 µg/mL streptomycin (all from Sigma, UK). 293F suspension cells were grown in Freestyle 293 expression medium (Life Technologies 12338-018) on a shaker incubator. Cells were grown at 37°C, 5% CO_2_ in a humidified incubator. Viruses were diluted and grown in Virus grown medium (VGM), which is DMEM with 0.1% bovine serum albumin (Sigma A0336), 10 mM HEPES, and glutamine, Penicillin and Streptomycin as in D10.

### Influenza Viruses and control sera

H1N1 viruses from the years 1977 - 2018 and H3N2 viruses were obtained from the Worldwide Influenza Centre at The Crick Institute (London, UK). Other reassortant viruses and control sheep sera were obtained from NIBSC, UK.

### Ethics and Study Approval

The study was in compliance with good clinical practice guidelines and the Declaration of Helsinki. The protocol was approved by the Research and Ethics Committee of Chang Gung Memorial Hospital, Beijing Ditan Hospital and the Weatherall Institute of Molecular Medicine. All subjects provided written informed consent. The list of donors with their details and isolated antibodies are included in Table 1.

### Isolation of human monoclonal antibodies

Antibodies were isolated from individual humans who either received seasonal influenza vaccine or were naturally infected with H7N9 virus in China or Taiwan. Antibodies were isolated using single cell isolation and cloning methods as described in detail previously (Tiller et al., 2008, Smith et al., 2009, Huang et al., 2015, Rijal et al., 2019). Briefly, plasmablasts in PBMC were stained (CD3neg, CD19pos, CD20lo/neg,CD27hi,CD38hi) and sorted as single cells using flow cytometry. mRNA from single plasmablasts was reverse transcribed to DNA and VH and Vk/λ genes were amplified using gene specific primers, then cloned into expression vectors containing IgG1 Heavy and Vk and Vλ constant regions. Heavy and light chain plasmids were co-transfected into 293T or ExpiCHO cells (Life Technologies A29133) for antibody expression.

### Antibody Screening

mAbs were initially screened for binding to MDCK-SIAT1 cells infected with either H1N1 or H3N2 viruses, and for lack of binding to HA protein expressed in stably transfected MDCK-SIAT1 cells. Binding to NA was confirmed by immuno-precipitation with infected cells or binding to 293T cells transfected with the NA gene of interest.

### Production of NA proteins

Tetrameric neuraminidase proteins were expressed from constructs based on the design of Xu et al. (Xu et al., 2008). In our version the signal sequence from A/PR/8/1934 HA was followed by the 15 residue tetramerization domain and thrombin site, followed by the NA sequence amino acids 69-469 (N1 numbering). Sequences were synthesised as human codon optimised cDNAs by Geneart and cloned into pCDNA3.1/-for transfection. HEK293F cells were transiently transfected using PEI-pro as a transfection reagent. Protein supernatant harvested 5-7 days post-transfection was titrated for NA activity in an ELLA and stored in aliquots at −80°C.

N9 NA protein (A/Anhui/1/13) was kindly provided by Donald Benton (The Francis Crick Institute) (Benton et al., 2017). The expression construct consisted of ectodomain residues 75-465 with and N-terminal 6x His tag, a human vasodilator-stimulated phosphoprotein tetramerization domain (Xu et al., 2008) and a TEV cleavage site under the control of promoter with a gp67 secretion signal peptide. The protein was expressed in Sf9 insect cells using a recombinant baculovirus system (Life Technologies). The protein was purified on a cobalt resin column and further purified by gel filtration to ensure the removal of monomeric and aggregated protein.

For antibody inhibition measurements a dilution of the NA containing supernatant was chosen that had just reached plateau activity in an ELLA. The sequences of all the constructs with their identification numbers are shown in Supplementary Table 1.

### NA inhibition assay: Enzyme-Linked Lectin Assay (ELLA)

ELLA assay was adapted from the methods described by Schulman et al. (Schulman et al., 1968) and Sandbulte et al. (Sandbulte et al., 2009). This assay detects the inhibition of NA enzymatic activity, cleavage of sialic acid, by anti-NA antibodies (Figure S3). Viruses or recombinant NA proteins were used as the source of NA. Virus growth medium was used to dilute antibodies and viruses. A Nunc Immunoassay ELISA plate (Thermo Scientific 439454) was plated overnight with 25 μg/ml fetuin (Sigma, F3385). Two-fold serial dilutions of sera or mAbs performed in duplicates were incubated together with a fixed amount of titrated NA source. Column 11 of a plate was used for NA source only control, and column 12 was used for medium only control. After 2 h incubation, antibody/NA mix were transferred to the PBS washed fetuin plate and incubated for 18-20 h at 37 °C buffered by CO_2_ as for tissue culture. Next day, the contents of the plate were discarded, and the plate washed 4 times with PBS. HRP conjugated peanut agglutinin (PNA-HRP, Sigma, L7759) at 1 μg/ml was added to the wells. PNA binds to the exposed galactose after cleavage of sialic acid by NA. After 1 h incubation and PBS wash, signal was developed by adding OPD (o-Phenylenediamine Dihydrochloride) solution (Sigma, P9187) and the reaction stopped after 5-15 min using 1 M H_2_SO_4_. Absorbance was read at 492 nm in a Clariostar plate reader (BMG Labtech).

### *In vivo* prophylaxis protection

All animal procedures were approved by an internal University of Oxford Ethics Committee and the United Kingdom Home Office. The experiments were carried out in accordance with the ‘Guide for the Care and Use of Laboratory Animals’, the recommendations of the Institute for Laboratory Animal Research, and Association for Assessment and Accreditation of Laboratory Animal Care International standards. Principle of the 3Rs were applied in design of experiments.

Mice used in protection studies, DBA/2OlaHsd mice (n=6/group) for X-179A and BALB/cOlaHsd (n=6/group) for PR8 viruses were purchased from Envigo, UK and housed in individually vented cages in a special unit for infectious diseases. Mice were anesthetised by isofluorane (Abott) and 50 μl of virus was administrated intranasally 24 hours after the intraperitoneal administration of 10 mg/kg antibody (500 μl). Mice were under regular observation and weighed. Mice with weight loss ≅20 percent or morbid clinical scores were euthanized by rising concentration of CO_2_. Non-specific IgG antibody was used as a negative control. Known HA-specific antibodies were used as positive controls. Mice were infected intranasally with lethal dose of viruses: X-179A (150 LD_50_, 10^4^ TCID_50_) and PR8 (1000 LD_50_, 10^4^ TCID_50_).

### Data and Statistical analysis

Graphs were generated using GraphPad Prism (version 9) and Microsoft Excel 2010. The ELLA titres were expressed as half maximal effective concentrations (EC_50_: midpoint between negative and plateau positive controls) derived by linear interpolation from neighbouring points in the titration curve. Kaplan Maier tests were performed to analyse the difference in mortality between experimental and control group mice. P values of <0.05 were considered as significant statistical difference.

## Acknowledgements

We acknowledge the flowcytometry services (Craig Waugh) provided by the Weatherall Institute of Molecular Medicine and the Core Instrument Center of Chang Gung University. We thank Donald Benton (The Francis Crick Institute) for providing N9 neuraminidase protein. These studies were funded by the Chinese Academy of Medical Sciences (CAMS) Innovation Fund for Medical Sciences (CIFMS), China (grant number: 2018-I2M-2-002), Townsend-Jeantet Prize Charitable Trust (Charity Number 1011770), Chang Gung Medical Research Program grant (CMRPG3G0921, CMRPG3G0922 and CORPG3J0111) and Ministry of Science and Technology of Taiwan (MOST 107-2321-B-182A-003-, MOST 108-2321-B-182A-001-). The work of the Crick Worldwide Influenza Centre, a WHO Collaborating Centre for Reference and Research on Influenza, was supported by the Francis Crick Institute receiving core funding from Cancer Research UK (FC001030), the Medical Research Council (FC001030) and the Wellcome Trust (FC001030). Views are those of the authors and do not necessarily reflect those of the funding bodies or employing institutes.

## Author Contributions

Conceptualisation: A.R.T., P.R., K.-Y.A.H.; Methodology: A.R.T., P.R., K.-Y.A.H; Investigation: P.R., A.R.T., K.-Y.A.H, B.B.Y., L.S., T.K.T., P.J., R.D.; Writing – Original Draft: P.R. and A.R.T.; Writing – Review & Editing: P.R., A.R.T., K.-Y.A.H., and R.S.D.; Supervision: A.R.T., K.-Y.A.H., P.R., R.S.D., J.W.M., and T.D.

## Supporting Information

**Table S1.**
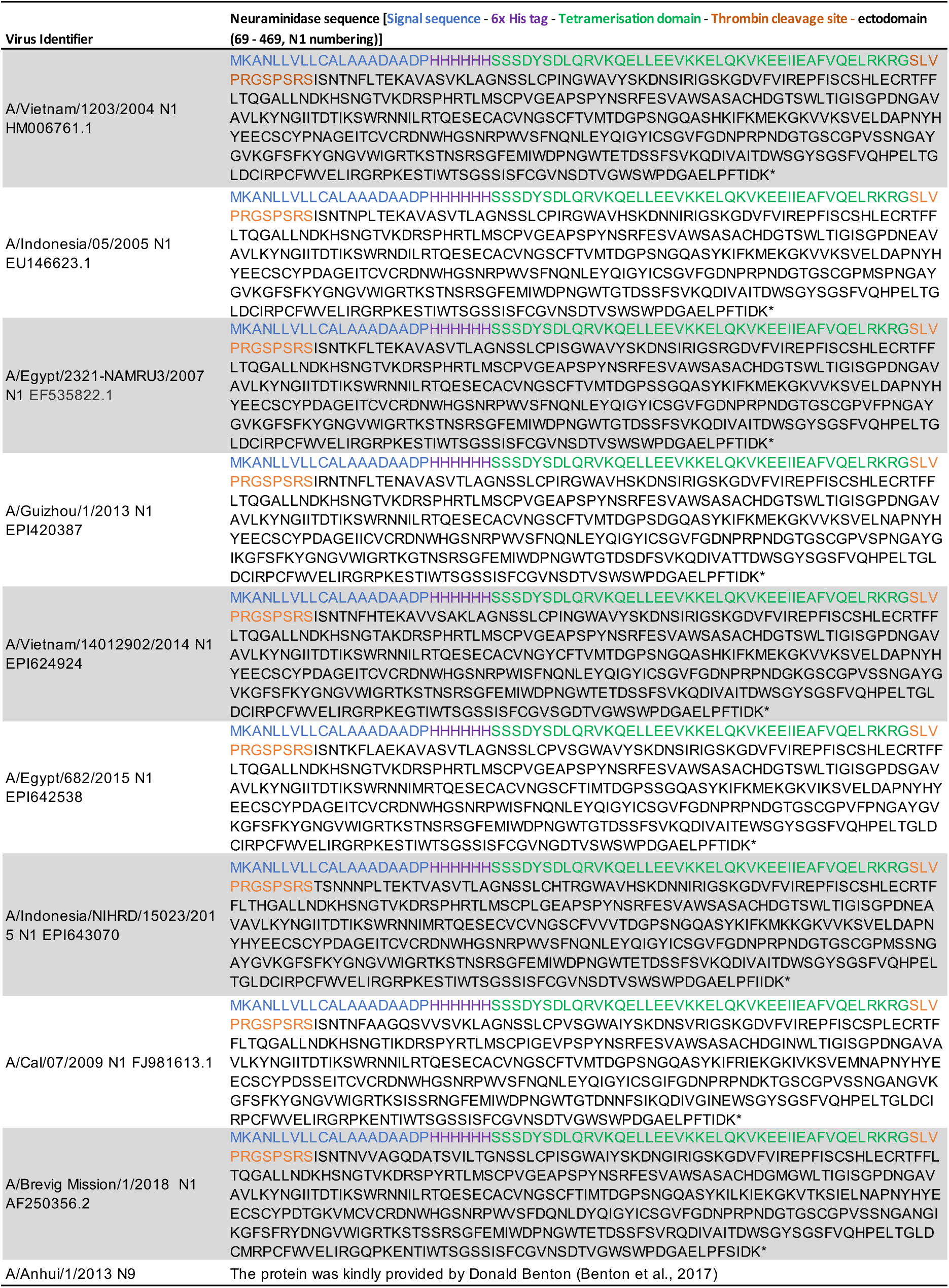
Sequences of secreted NA proteins

**Figure S1.**
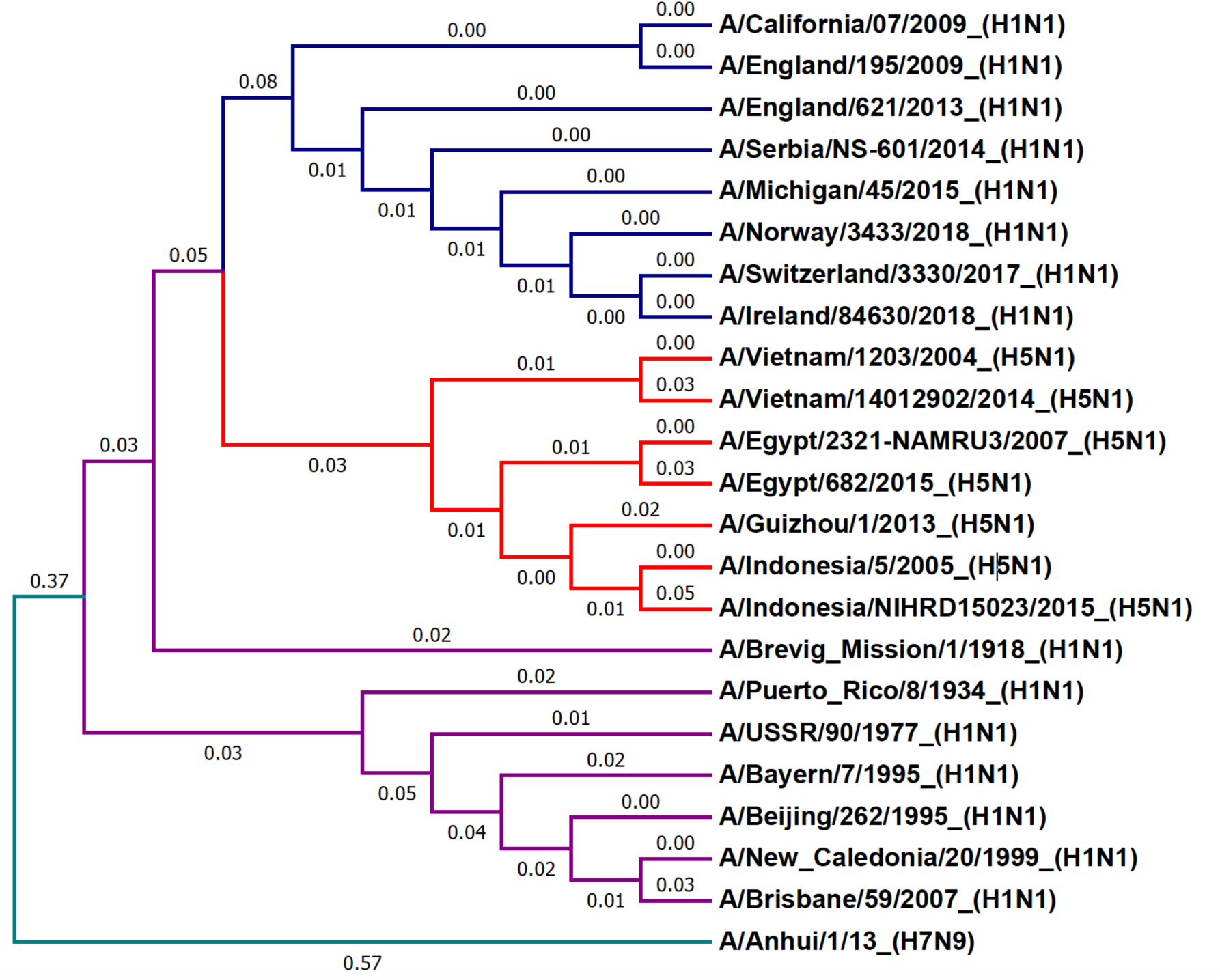
The phylogenetic tree of N1 and N9 neuraminidases used in this paper. The values on branches shows the evolutionary distances between neuraminidases. Made using MEGA7 software, muscle alignment and neighbour-joining tree settings.

**Figure S2.**
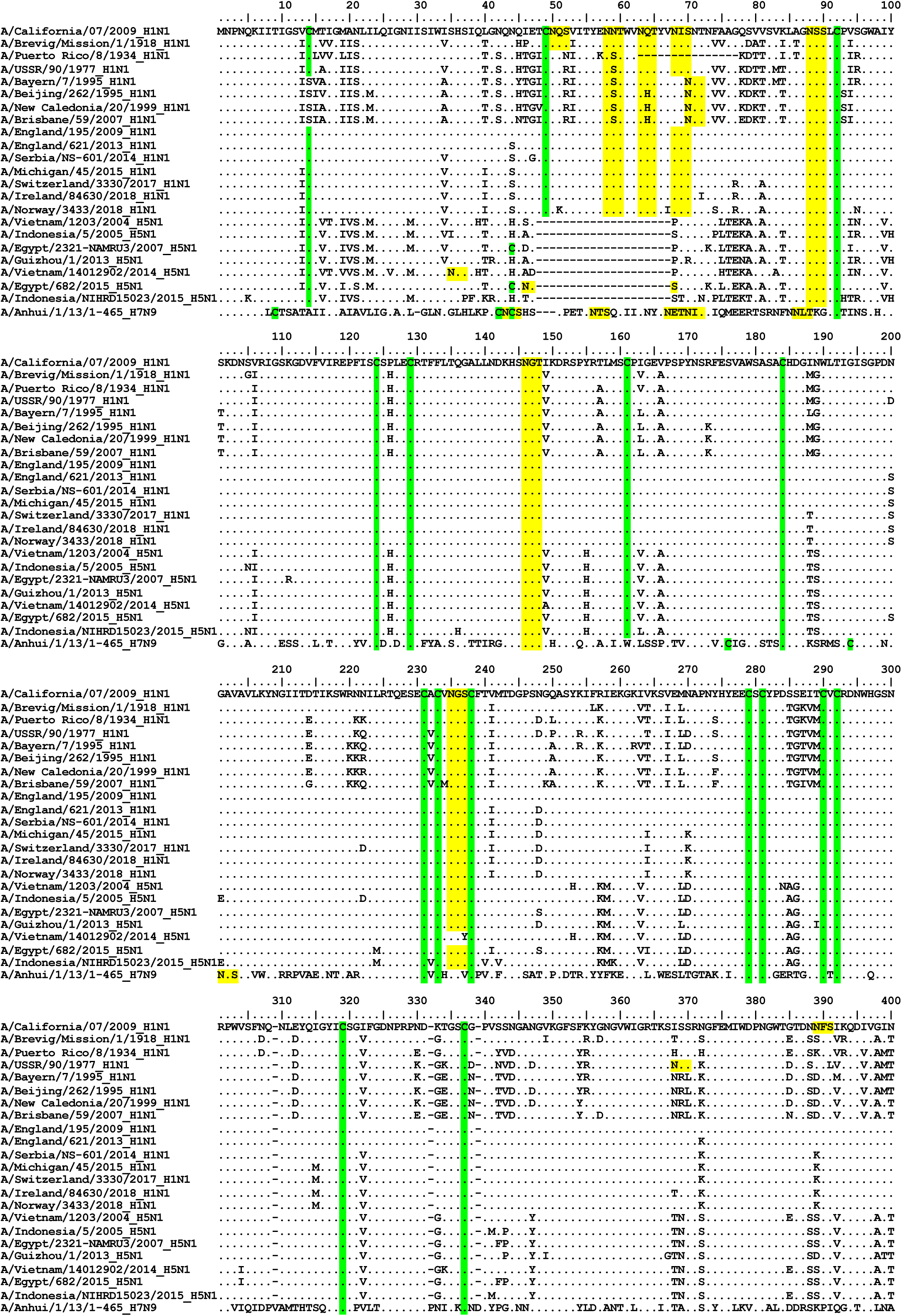
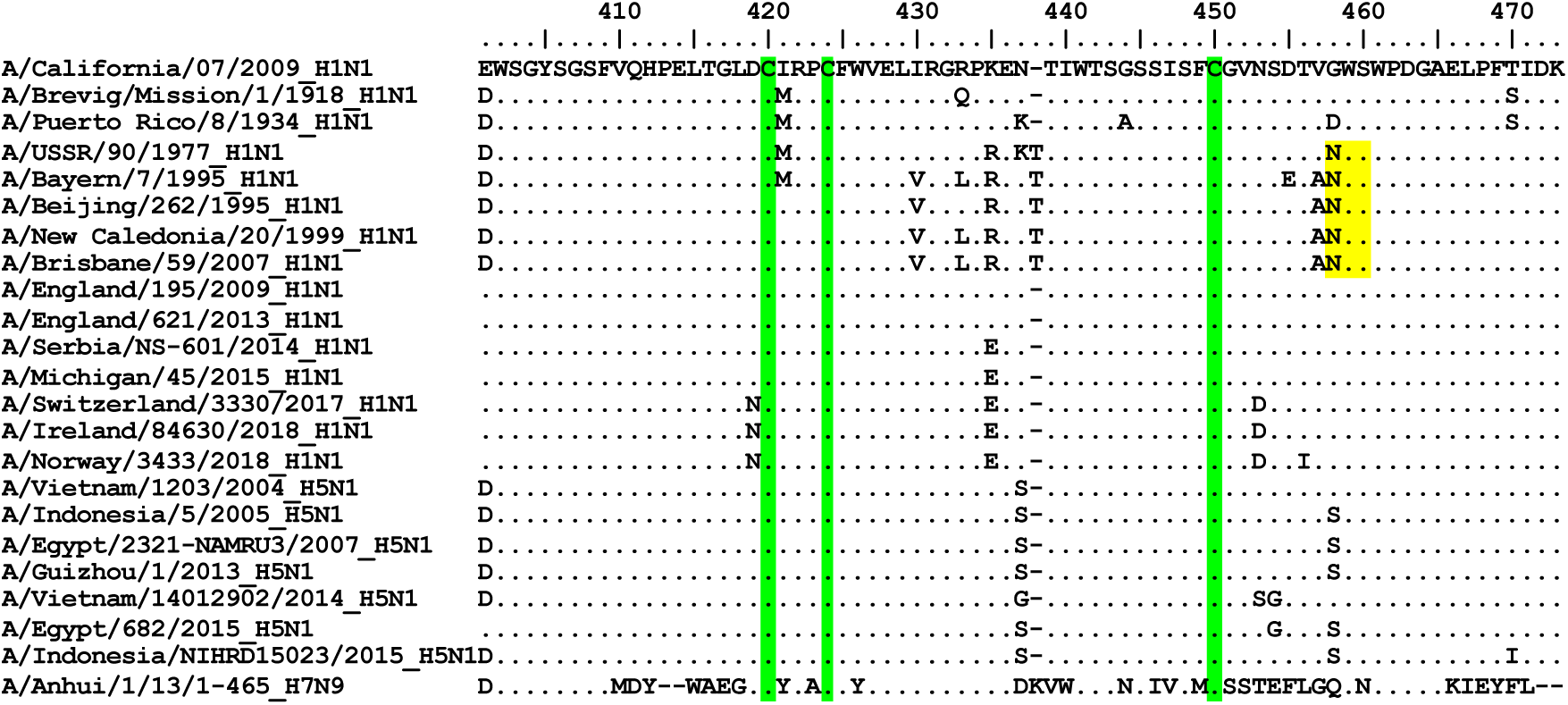
Amino acid sequence alignment of the neuraminidases used in this paper. The numbering is not NA numbering and is only for alignment purpose. The alignment was done using BioEdit software.

**Figure S3.**
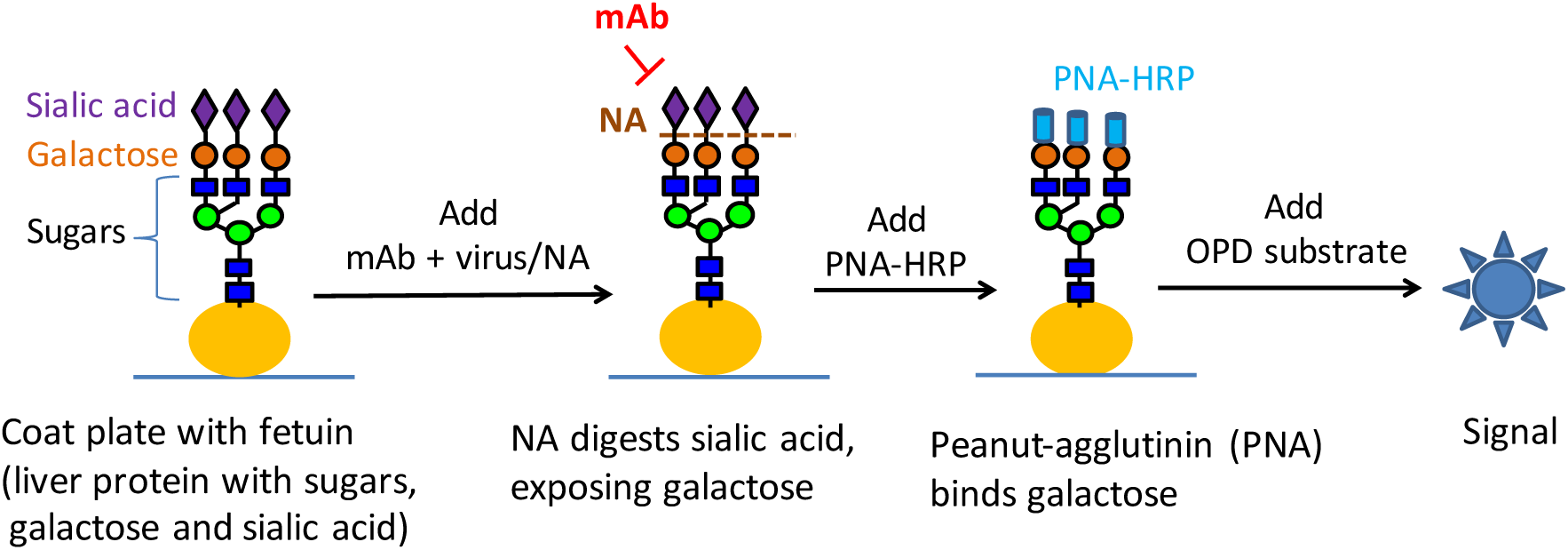
Schematic figure of ELLA assay

